# KINOMO: A non-negative matrix factorization framework for recovering intra- and inter-tumoral heterogeneity from single-cell RNA-seq data

**DOI:** 10.1101/2022.05.02.490362

**Authors:** Somnath Tagore, Yiping Wang, Jana Biermann, Raul Rabadan, Elham Azizi, Benjamin Izar

**Affiliations:** Department of Systems Biology, Columbia University Irving Medical Center, New York, NY, USA; Department of Medicine, Division of Hematology/Oncology, Columbia University Irving Medical Center, New York, NY, USA; Program for Mathematical Genomics, Columbia University, New York, NY, USA; Columbia Center for Translational Immunology, New York, NY, USA; Department of Biomedical Informatics, Columbia University, New York, NY, USA; Department of Biomedical Engineering, Columbia University, New York, NY, USA; Irving Institute for Cancer Dynamics, Columbia University, New York, NY, USA

**Keywords:** Dimensionality reduction, intra-tumor heterogeneity, inter-tumor heterogeneity, non-negative matrix factorization, single-cell RNA-seq

## Abstract

Single-cell RNA-sequencing (scRNA-seq) is a powerful technology to uncover cellular heterogeneity in tumor ecosystems. Due to differences in underlying gene load, direct comparison between patient samples is challenging, and this is further complicated by the sparsity of data matrices in scRNA-seq. Here, we present a factorization method called KINOMO (**K**ernel d**I**fferentiability correlation-based **NO**n-negative **M**atrix factorization algorithm using Kullback-Leibler divergence loss **O**ptimization). This tool uses quadratic approximation approach for error correction and an iterative multiplicative approach, which improves the quality assessment of NMF-identified factorization, while mitigating biases introduced by inter-patient genomic variability. We benchmarked this new approach against nine different methods across 15 scRNA-seq experiments and find that KINOMO outperforms prior methods when evaluated with an adjusted Rand index (ARI), ranging 0.82-0.91 compared to 0.68-0.77. Thus, KINOMO provides an improved approach for determining coherent transcriptional programs (and meta-programs) from scRNA-seq data of cancer tissues, enabling comparison of patients with variable genomic backgrounds.

**Classification:** Physical Sciences (Applied Mathematics; Biophysics and Computational Biology), Biological Sciences (Applied Biological Sciences; Biophysics and Computational Biology; Medical Sciences; Systems Biology.).

**Significance Statement:** Identification of shared or distinct cell programs in single-cell RNA-seq data of patient cancer cells is challenging due to underlying variability of gene load which determines transcriptional output. We developed an analytical approach to define transcriptional variability more accurately across patients and therefore enable comparison of program expression despite inherent genetic heterogeneity. Thus, this method overcomes challenges not adequately addressed by other methods broadly used for the analysis of single-cell genomics data.

## Introduction

Non-negative matrix factorization (NMF) is a widely used method for performing dimensionality reduction and feature extraction on ‘non-negative’ data **(Lee et al, 1999)**. The major difference between NMF and other factorization methods such as Singular-value decomposition (SVD) is that it performs feature extraction on non-negativity data thus allowing additive combinations of intrinsic features. Likewise, most of the low-rank matrix dimensionality reduction mechanisms are unsupervised methods that are capable of identifying the low-dimensional structure embedded in the original data. A non-negativity constraint particularly helps in interpreting big and complex data structure, thereby preserving physical feasibility. In the field of next-generation sequencing, such as single-cell RNA-sequencing (scRNA-Seq) which involves non-negative measurements of gene expression, emergence of these mathematical techniques could help in better understanding of the underlying biological processes. scRNA-seq is a powerful method to identify cell identity (e.g., hematopoietic lineage) and activity programs (e.g., cell cycle, differentiation states) across non-transformed lineages **(Kotliar et al, 2019)**.

In scRNA-seq studies of patient cancer tissues, the most important driver for inter-patient heterogeneity among malignant cells is the patient of origin **(Patel et al, 2014; Tirosh et al, 2016)**. These studies highlighted that gene load (e.g., oncogenic mutations or aneuploidy patterns) were the key determinant of transcriptional output, thus, posing a major bias when comparing gene expression between two patients and among potential cancer cell subpopulations within the same patient with varying genomic patterns. Thus, while intra-patient transcriptional variability may inform important processes (e.g., metastatic behavior, drug response/resistance) **(Jerby-Arnon et al, 2018)** and dissecting heterogeneity and cell states across patient populations can be highly informative, direct comparison between patients is challenging. For this purpose, factorization approaches provide an immediate benefit for understanding this heterogeneity, by denoising the sparse signal of scRNA-seq data.

Conventional NMF **(Lee et al, 1999)** decomposes a matrix *A* into two matrices with non-negative entries with smaller factors, *A* ≈ *WH*, where *A* ∈ ℝ^*n×m*^, *W* ∈ ℝ^*n×k*^, *H* ∈ ℝ^*k×m*^ Without loss of generalization, rows of *A* represent features (e.g., genes) and columns of *A* represent samples. Depending on context, *W* can be interpreted as a feature mapping. Rows of *W* represent disease profiles or metagenes **(Brunet et al, 2004)**. Columns *H* are compact representations of samples, i.e., sample profiles. Various NMF methods have been published to date focusing on different application domains, such as sparse NMF (SNMF), discriminant NMF (DNMF) for RNA-seq data **(Jia et al., 2015; Kim & Park, 2007)**, among others. SNMF, for example, introduces a regularization term on *W* or *H* to control the degree of sparsity and generate sparser representation, whereas DNMF incorporates Fisher’s discriminant criterion in the coefficient matrix by maximizing the distance among any samples from different classes meanwhile minimizing the dispersion between any pair of samples in the same class. The major problem in scRNA-seq data is the difficulty in assigning specific classes to cells, since this information is not known confidently. Moreover, most of the scRNA-seq methodologies suffer from various technical issues like amplification bias, differences in library sizes, dropouts, among others **(Buettner et al., 2015; Kharchenko, Silberstein & Scadden, 2014)**.

In this paper, we propose **K**ernel d**I**fferentiability correlation-based **NO**n-negative **M**atrix factorization algorithm using Kullback-Leibler divergence loss **O**ptimization (KINOMO), a semi-supervised NMF model that is robust to noise and also uses ‘prior’ biological knowledge for better refinement. We benchmark our approach against nine frequently used NMF approaches and demonstrate that KINOMO outperforms these across multiple scRNA-seq data sets measured by adjusted Rand index (ARI). Thus, KINOMO enables more accurate identification and comparison of cell states tissues and overcomes critical challenges in the analysis of scRNA-seq data that includes cancer cells with variable gene load. KINOMO is freely available for research use on GitHub (https://github.com/IzarLab/KINOMO).

## RESULTS

### KINOMO

KINOMO has three major steps, including 1) filtering and cleaning, 2) NMF core module, and 3) metagene and factor block estimation. The filtering and cleaning steps begin with the tumor cell estimation of an individual scRNA-seq sample, followed by normalization and scaling. The second step consists of two sub-steps a) Factorization Error Analysis, using L_2,1_ norm loss to handle outliers, adding prior knowledge by introducing graph regularization parameters, sequential quadratic approximation for Kullback-Leibler divergence loss, local geometrical structure preservation, optimizing the update rules for approximation matrix *WH* and clustering using Kernel differentiability correlation; and b) factor factor survey analysis. The third step consists of metagene and factor block estimation.

### Factorization Error Analysis

For a non-negative matrix *X*^*mXn*^, denoting single cell data with *m* features (e.g. gene expression) measured for *n* cells, KINOMO can decompose it into two non-negative matrices *W*^*mXk*^ and *H*^*kXn*^, such that *X ≈ H*, where *k < min (m,n)*. The Euclidean distance between *X* and its approximation matrix *WH* is applied to minimize the factorization error, which can be written as **(Lee and Seung, 2001)**,

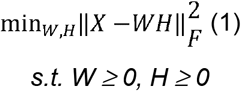

Here, 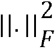 is the Frobenius norm of a matrix. Since, it is difficult to find a global minimal solution by optimizing the convex non-linear objective function, KINOMO adopts the multiplicative iterative update algorithm to approximate *W* and *H*,

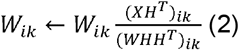

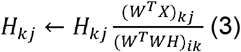

For the multiplicative iterative approach that KINOMO considers, *W* and *H* are initialized randomly, and in each repetition, the update steps are processed until maximum number of iterations are reached. The stopping criteria if fulfilled when the shift between two iterations is negligible. Moreover, the update steps are based on the mean squared error objective function. This approach is different from other multiplicative methods such as NMFNA **(Ding et al, 2021)**, which relies on Karush–Kuhn–Tucher conditions.

### Using L_2,1_ norm loss to handle outliers

As the representation above **(Lee and Seung, 2001)** fails to handle outliers and noise, **Kong et al., 2011** replaced the Frobenius norm with L_2,1_ norm loss as defined below,

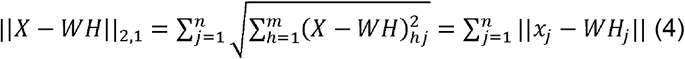

Here, the error of each data point is ||*x*_*j*_ − *WH*_*j*_|| rather than squared, *j*^*th*^ column of *X*, thus the errors caused by outliers and noise do not dominate the objective function compared to the L2 norm.

### Adding scRNA-seq derived prior knowledge by introducing graph regularization parameters

**Chen and Zhang, 2018** proposed a method of introducing an error matrix *S* ∈ ℝ^*n×m*^. For a sample with multiple modalities (for the purpose of explanation, we consider the number of modalities = 2), defined as two variables 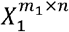 and 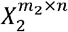 with *m*_*1*_ and *m*_*2*_ features respectively, measured for *n* cells, Chen and Zhang defined the optimization problem as,

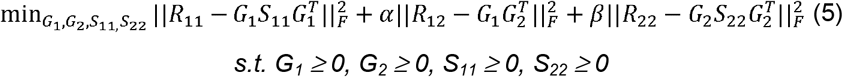

Here 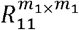 and 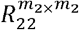 are the symmetric feature similarity matrices of *X*_*1*_ and *X*_*2*_, respectively, that is, their respective co-expression networks; 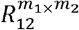 is the two-type feature similarity matrix between *X*_1_ and 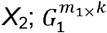 and 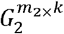 are the non-negative factored matrices used for identifying modules in their respective networks; 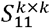 and 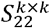 are also symmetric matrices whose diagonal elements can be used for measuring associations between identified modules; *k* is the user prespecified dimension parameter; α and β are graph regularization parameters in the objective function and default settings are *m*_1_/*m*_2_ and (*m*_1_/*m*_2_)^2^, respectively **(Chen and Zhang, 2018)**. While classical non-negative factorization approaches fit the data in Euclidean space, KINOMO uses the intrinsic geometry of scRNA-seq and incorporates it as additional regularization terms *(α* and *β*).

### Sequential quadratic approximation for Kullback-Leibler divergence loss

The error function is a suitably chosen function. Here, we chose the Kullback-Leibler divergence **(Kullback-Leibler, 1951)**. Thus, assuming *W* is known and *H* is to be solved. Let,

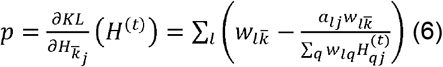

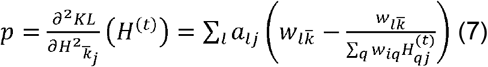

Here, *H*^(*t*)^is the current value of *H* in the iterative procedure and 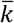 is mean factor. When fixing all the other entries, the Taylor expansion of the penalized KL divergence up to the 2^nd^ order at 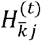 w.r.t. 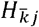 is

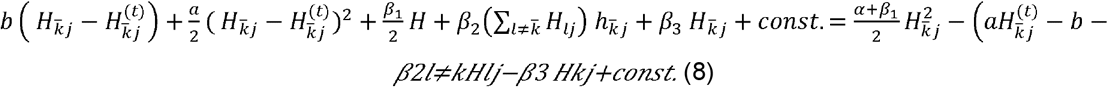

This can be solved explicitly by

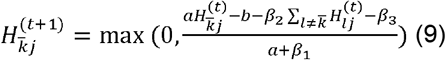

Similarly, we update 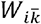.

Simplifying the above term, we have

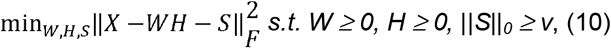

where *v* specifies the maximum number of non-zero elements in *S*. Likewise,

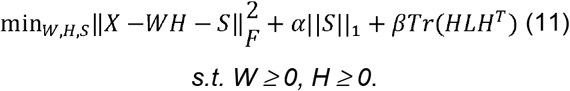

The graph regularized constraint indicates the inherent geometrical structure of the input networks. In other words, the graph regularized constraint ensures that interactive features in the Euclidean space are also close to each other in the low-dimensional space. We can construct a weight matrix µ, by using the markers gene of cell groups (as an example) using heat kernel weighting between *x*_*i*_ and *x*_*j*_:

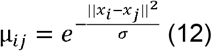

Here, *σ* is parameter that controls the weighting and we set *σ = 1* in the experiment. The weight matrix is constructed by a predefined set of marker genes, which for example in melanoma, such as markers of melanocytic lineage (*MITF, PMEL, TYR*), neuronal development and differentiation, (*NGFR, NLGN3, NRXN*), and synapse function and formation (e.g., *SNCA, SYT11, GPHN*).

### Local geometrical structure preservation

We use Euclidean distance to measure the dissimilarity in the low-dimensional representation of *X*:

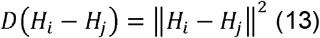

The local geometrical structure preserving criterion is:

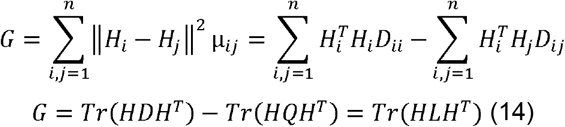

Here, *D* = ∑_*j*_*Q*_*ij*_ is a diagonal matrix, *L* = *D* – *Q* is graph Laplacian. Thus, minimizing G will result in *H*_*i*_ and *H*_*j*_ to be close to each other, if data point *x*_*i*_ and *x*_*j*_ are close, thus the distance relation between points in *X* is preserved in low dimensional matrix *H* **(Zhai et al., 2020)**.

### Optimizing the update rules for approximation matrix WH

The optimization function of KINOMO can be rewritten as:

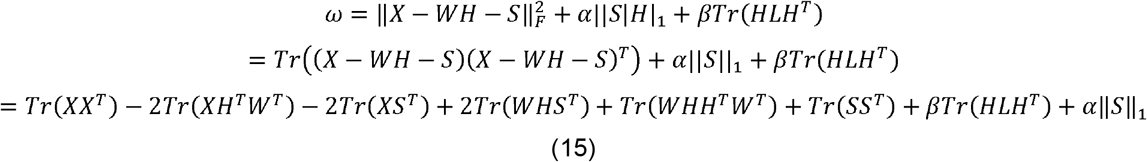

The Lagrange function *L* is stated as

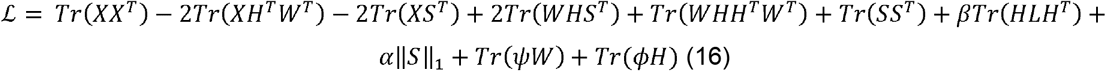

Here *ψ, ϕ* are Lagrange multipliers.

The partial derivatives of ℒ with respect to *W* and *H* are:

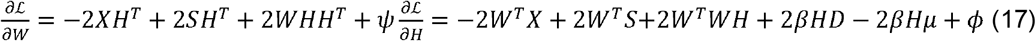

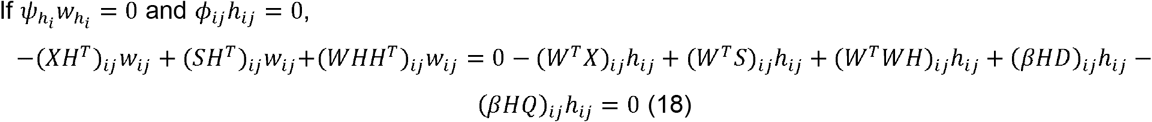

Updating rules for *W* and *H* are:

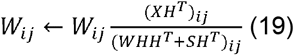

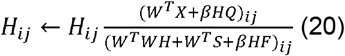

### Clustering using Kernel differentiability correlation

We define a continuous version of differentiability for each cell *i* in gene *k*

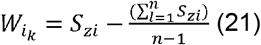

We now calculate the differentiability correlation among the given *n* cells in the data set

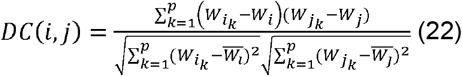

Here,

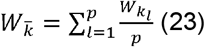

The Kernel DC is constructed as

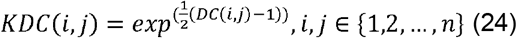

### Rank factor analysis, metagene selection and co-correlation analysis

A critical parameter in KINOMO is the factorization factor, *r*. It defines the number of metagenes used to approximate the target matrix. Given an NMF method and the target matrix, a common way of deciding on *r* is to optimize according to a quality measure of the results scanning a range of *r* values. **Brunet et al, 2004** proposed to take the first value of *r* for which the cophenetic coefficient starts decreasing, whereas, **Hutchins et al, 2008** suggested to choose the first value where the Residual Sum of Squares (RSS) curve presents an inflection point, and **Frigyesi et al, 2008** considered the smallest value at which the decrease in the RSS is lower than the decrease of the RSS obtained from random data. KINOMO selects the optimal factor by selecting the first- and second-best value of *r* for which the cophenetic coefficient starts decreasing. For each best and second-best factor factor (per sample), the top 30/50/100/200/300 top metagenes are selected using the KINOMO score. A co-correlation analysis using Spearman’s correlation is performed for identifying the correlations among all factors across all samples (best and second-best factor factor done separately). Finally, the consensus factor factors are selected using significance testing (Kruskal-Wallis p-value) and/or correlation value among all factors with a correlation threshold of 0.4–0.9. This threshold range has been selected by running KINOMO on multiple datasets. This is done iteratively, unless consensus factor blocks are obtained.

### Description of the datasets

To determine the performance of our method, we selected 15 samples from three publicly available scRNA-seq studies **(Table 1)**. For the purpose of validating our method, we only considered the tumor compartments from these 15 samples. *First*, a non-small cell lung cancer (NSCLC) dataset consisting in total of 42 tissue biopsy samples from stage III/IV NSCLC patients by scRNA-seq **(Wu et al, 2021)**. Five out of 42 samples were selected based on higher median genes per cells. The estimated number of cells in these 5 samples (GSM4453576_P1, GSM4453578_P3, GSM4453584_P9, GSM4453592_P17, GSM4453616_P41) ranged from 3,521–8,700, whereas the median genes per cell ranged from 1,566– 2,220, respectively. Subsequently, we predicted the tumor cells from all these individual samples based on inferring copy-number alterations using *inferCNV* **(Tickle et al, 2019)**. The number of inferred tumor cells in these five samples ranged from 2,194–8,585, respectively. *Second*, a prostate cancer (PCa) dataset consisting of transrectal prostate biopsies (n = 3) and radical prostatectomy (RP) specimens (n = 8), half of which had matched benign-appearing tissue **(Song et al, 2022)**. Again, we selected five samples (PA_AUG_PB_1A_S1, PA_PB1A_Pool_1_3_S50_L002, PA_PB1B_Pool_1_2_S74_L003, PA_PR5186_Pool_1_2_3_S27_L001, PA_PR5269_1_S25_L002) with the estimated number of cells ranging from 766–1,148, whereas median genes per cell ranging from 1,205–1,949. Further, the number of tumor cells in these five samples ranged from 277–630, respectively. *Third*, an osteosarcoma (OS) dataset consisting of RNA-sequencing of 100,987 individual cells from 7 primary, 2 recurrent, and 2 lung metastatic osteosarcoma lesions **(Zhou et al, 2020)**. In this data set, for the five selected samples (GSE152048_BC10, GSE152048_BC11, GSE152048_BC16, GSE152048_BC2, GSE152048_BC20) the estimated number of cells ranged from 5,900–17,000, whereas and median genes per cell ranged from 1,000–1,900. The number of tumor cells in these five samples ranged from 1,700–13,000, respectively.

**Table 1.** Overview of the datasets used in the study.

| Dataset | Cancer cohort | Study | Estimated Number of Cells | Median Genes per Cell | Number of tumor cells |
| --- | --- | --- | --- | --- | --- |
| GSM4453576_P1 | Non-small cell lung cancer | Wu et al, 2021 | 5,320 | 2,220 | 3,822 |
| GSM4453578_P3 | Non-small cell lung cancer | Wu et al, 2021 | 7,835 | 1,632 | 7,668 |
| GSM4453584_P9 | Non-small cell lung cancer | Wu et al, 2021 | 3,521 | 1,860 | 2,194 |
| GSM4453592_P17 | Non-small cell lung cancer | Wu et al, 2021 | 8,700 | 1,566 | 8,585 |
| GSM4453616_P41 | Non-small cell lung cancer | Wu et al, 2021 | 6,222 | 1,863 | 5,676 |
| PA_AUG_PB_1A_S1 | Prostate cancer | Song et al, 2022 | 766 | 1,205 | 277 |
| PA_PB1A_Pool_1_3_S50_L002 | Prostate cancer | Song et al, 2022 | 952 | 1,301 | 466 |
| PA_PB1B_Pool_1_2_S74_L003 | Prostate cancer | Song et al, 2022 | 946 | 1,439 | 430 |
| PA_PR5186_Pool_1_2_3_S27_L001 | Prostate cancer | Song et al, 2022 | 854 | 1,949 | 479 |
| PA_PR5269_1_S25_L002 | Prostate cancer | Song et al, 2022 | 1,148 | 1,602 | 630 |
| GSE152048_BC10 | Osteosarcoma | Zhou et al, 2020 | 17,481 | 1,915 | 13,460 |
| GSE152048_BC11 | Osteosarcoma | Zhou et al, 2020 | 13,444 | 1,899 | 11,024 |
| GSE152048_BC16 | Osteosarcoma | Zhou et al, 2020 | 10,210 | 1,964 | 3,063 |
| GSE152048_BC2 | Osteosarcoma | Zhou et al, 2020 | 5,962 | 1,002 | 1,789 |
| GSE152048_BC20 | Osteosarcoma | Zhou et al, 2020 | 11,096 | 1,988 | 9,210 |

### Performance evaluation

To evaluate how well the inferred clusters recovered the true subpopulations in the scRNA-seq data, we used the Hubert-Arabie Adjusted Rand Index (ARI) for comparing two partitions **(Rand 1971)**. The metric is adjusted for chance, such that independent clustering has an expected index of 0 and identical partitions have an ARI equal to 1, and was calculated using the implementation in the ‘mclust’ R package v5.4 **(Scrucca et al, 2016)**. We also used the ARI to evaluate the stability of the clusters, by comparing the partitions from each pair of the five independent runs for each method with a given number of clusters.

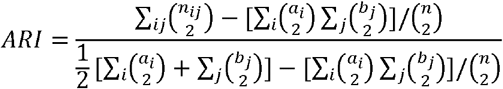

Here, *n* _*i,j*_ are values from the contingency table *a*_*i*_ and *b*_*j*_ denote the summation of the *i*^th^ row *j*^th^ column of the contingency table respectively.

To evaluate the similarities between the partitions obtained by different methods **(Table 2)**, we first calculated a consensus partition from the five independent runs for each method, using the ‘clue’ R package v0.3-55 **(Hornik 2005)**. Next, for each sample and each imposed number of clusters, we calculated the ARI between the partitions for each pair of methods, and used hierarchical clustering based on the median of these ARI values across all samples to generate a dendrogram representing the similarity among the clusters obtained by different methods. To investigate how representative this dendrogram is, we also clustered the methods based on each sample separately, and calculated the fraction of such dendrograms in which each subcluster in the overall dendrogram appeared. Finally, we investigated whether clustering performance was improved by combining two methods into an ensemble. For each sample, and with the true number of clusters imposed, we calculated a consensus partition for each pair of methods, and used the ARI to evaluate the agreement with the true cell labels. We then compared the ensemble performance to the performances of the two individual methods used to construct the ensemble. **Figure 1a-c** illustrates the ARI heatmap for evaluating the class assignment using KINOMO against all other methods for PCa, OS and NSCLC samples. For all the three sample sets, the result illustrates that KINOMO outperforms all the other methods in terms of better class assignment based on ARI score (0.91 in PCa, 0.87 in OS, 0.82 in NSCLC). The closest performing methods are NMF (0.77 in PCa, 0.75 in OS, 0.71 in NSCLC) and SC3 (0.74 in PCa, 0.72 in OS, 0.68 in NSCLC) respectively.

**Table 2.** Overview of the methods used in the study for benchmarking KINOME.

| Methods | Description | Reference |
| --- | --- | --- |
| Hierarchical_Clustering | Groups data over a variety of scales by creating a cluster tree or dendrogram | Maimon et al, 2006 |
| K-Means | Starts with a first group of randomly selected centroids, used as the beginning points for every cluster and then performs iterative calculations to optimize the positions of the centroids | Hartigan et al, 1979 |
| Fuzzy_C-Means | This algorithm works by assigning membership to each data point corresponding to each cluster center on the basis of distance between the cluster center and the data point. | Dunn 1973 |
| SC3 (v1.8.0) | PCA dimension reduction or Laplacian graph. K-means clustering on different dimensions. Hierarchical clustering on consensus matrix obtained by K-means | Strehl et al, 2003 |
| TSCAN | PCA dimension reduction followed by model-based clustering | Zhicheng et al, 2016 |
| SAFE (v2.1.0) | Ensemble clustering using SC3, CIDR, Seurat and t-SNE + Kmeans | Yuchen et al, 2019 |
| FlowSOM (v1.12.0) | PCA dimension reduction (dim=30) followed by self-organizing maps (5x5, 8x8 or 15x15 grid, depending on the number of cells in the data set) and hierarchical consensus meta-clustering to merge clusters | Van Gassen et al, 2015 |
| CIDR (v0.1.5) | PCA dimension reduction based on zero-imputed similarities, followed by hierarchical clustering | Lin et al, 2017 |
| NMF | Non-negative matrix factorization | Lee et al, 1999 |

**Figure 1.**
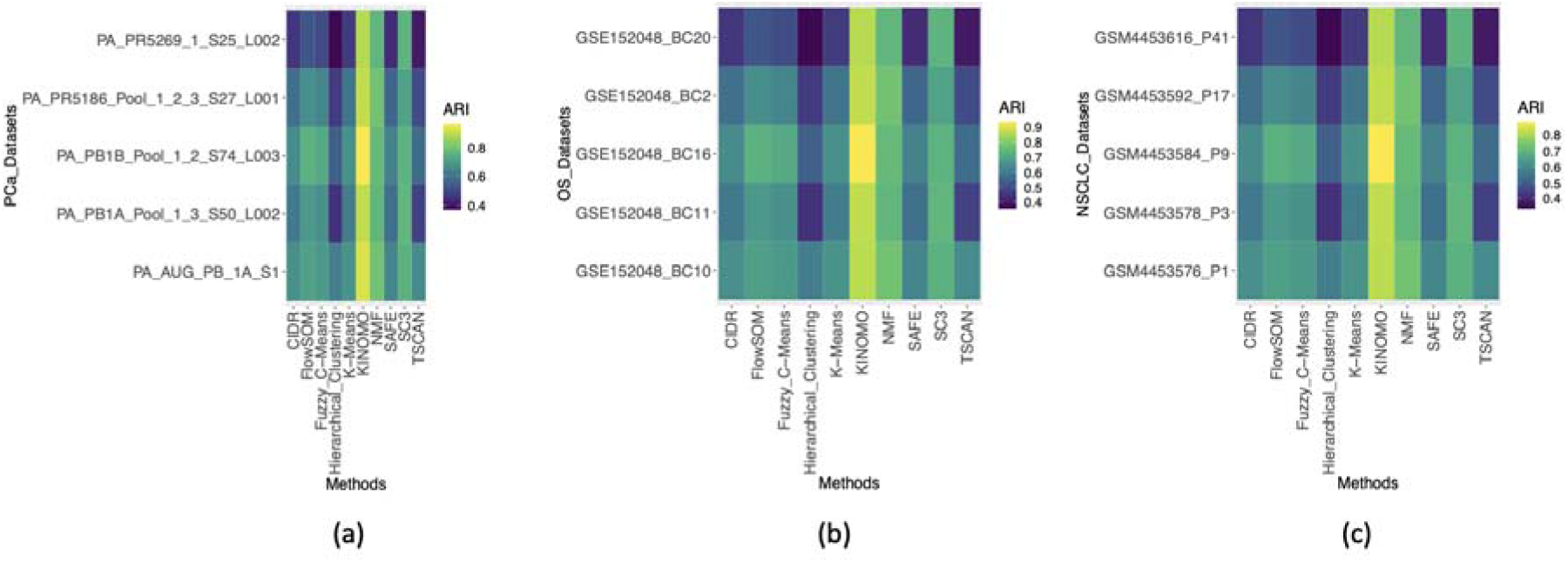
Adjusted Rand index (ARI) heatmap partition evaluation, for (a) PCa (KINOMO- 0.91), (b) OS (KINOMO- 0.91), and (c) NSCLC (KINOMO- 0.82) datasets.

**Tables 3-5** show the clustering results of the 15 samples. The final results are evaluated by taking the average value of ARI over 100 runs. Compared with conventional NMF, KINOMO shows increased performance over all the datasets. Moreover, the overall ARI for all methods on these 15 samples ranges between 0.3–0.8. The worst performing method is hierarchical clustering, which is expected due to the sparsity of single-cell data and it is difficult to have a correct exact prior knowledge about cell types. Thus, the gene expression programs are quite mixed across all clusters. Other well-performing methods apart from KINOMO are NMF, SC3 and Fuzzy C-Means **(Tables 3-5)**.

**Table 3.** Benchmarking KINOMO with other methods (ARI) in NSCLC dataset.

| Dataset | Hierarchical_Clustering | K-Means | Fuzzy_C-Means | SC3 | TSCAN | SAFE | FlowSOM | CIDR | NMF | KINOME |
| --- | --- | --- | --- | --- | --- | --- | --- | --- | --- | --- |
| GSM4453576_P1 | 0.572 | 0.621 | 0.633 | 0.691 | 0.602 | 0.591 | 0.653 | 0.611 | 0.731 | 0.816 |
| GSM4453578_P3 | 0.413 | 0.571 | 0.612 | 0.701 | 0.443 | 0.541 | 0.632 | 0.561 | 0.703 | 0.821 |
| GSM4453584_P9 | 0.501 | 0.613 | 0.671 | 0.692 | 0.531 | 0.583 | 0.691 | 0.603 | 0.711 | 0.877 |
| GSM4453592_P17 | 0.431 | 0.551 | 0.589 | 0.656 | 0.461 | 0.521 | 0.609 | 0.541 | 0.731 | 0.823 |
| GSM4453616_P41 | 0.342 | 0.433 | 0.462 | 0.696 | 0.372 | 0.403 | 0.482 | 0.423 | 0.701 | 0.811 |

**Table 4.** Benchmarking KINOMO with other methods (ARI) in PCa dataset.

| Dataset | Hierarchical_Clustering | K-Means | Fuzzy_C-Means | SC3 | TSCAN | SAFE | FlowSOM | CIDR | NMF | KINOME |
| --- | --- | --- | --- | --- | --- | --- | --- | --- | --- | --- |
| GSM4453576_P1 | 0.62348 | 0.67689 | 0.68997 | 0.75319 | 0.65618 | 0.64419 | 0.71177 | 0.66599 | 0.79679 | 0.92214 |
| GSM4453578_P3 | 0.45017 | 0.62239 | 0.66708 | 0.76409 | 0.48287 | 0.58969 | 0.68888 | 0.61149 | 0.76627 | 0.89489 |
| GSM4453584_P9 | 0.54609 | 0.66817 | 0.73139 | 0.75428 | 0.57879 | 0.63547 | 0.75319 | 0.65727 | 0.77499 | 0.95593 |
| GSM4453592_P17 | 0.46979 | 0.60059 | 0.64201 | 0.71504 | 0.50249 | 0.56789 | 0.66381 | 0.58969 | 0.79679 | 0.89707 |
| GSM4453616_P41 | 0.37278 | 0.47197 | 0.50358 | 0.75864 | 0.40548 | 0.43927 | 0.52538 | 0.46107 | 0.76409 | 0.88399 |

**Table 5.**
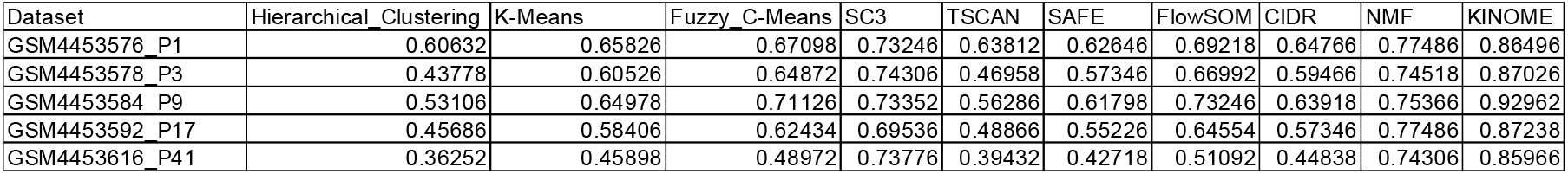
Benchmarking KINOMO with other methods (ARI) in OS dataset.

| Dataset | Hierarchical_Clustering | K-Means | Fuzzy_C-Means | SC3 | TSCAN | SAFE | FlowSOM | CIDR | NMF | KINOME |
| --- | --- | --- | --- | --- | --- | --- | --- | --- | --- | --- |
| GSM4453576_P1 | 0.60632 | 0.65826 | 0.67098 | 0.73246 | 0.63812 | 0.62646 | 0.69218 | 0.64766 | 0.77486 | 0.86496 |
| GSM4453578_P3 | 0.43778 | 0.60526 | 0.64872 | 0.74306 | 0.46958 | 0.57346 | 0.66992 | 0.59466 | 0.74518 | 0.87026 |
| GSM4453584_P9 | 0.53106 | 0.64978 | 0.71126 | 0.73352 | 0.56286 | 0.61798 | 0.73246 | 0.63918 | 0.75366 | 0.92962 |
| GSM4453592_P17 | 0.45686 | 0.58406 | 0.62434 | 0.69536 | 0.48866 | 0.55226 | 0.64554 | 0.57346 | 0.77486 | 0.87238 |
| GSM4453616_P41 | 0.36252 | 0.45898 | 0.48972 | 0.73776 | 0.39432 | 0.42718 | 0.51092 | 0.44838 | 0.74306 | 0.85966 |

### Application to non-small cell lung cancer (NSCLC) dataset

For the purpose of highlighting the features of KINOMO, we focus on the NSCLC dataset **(Wu et al, 2021)** consisting of biopsy samples from 42 advanced NSCLC patients with diverse histological and molecular phenotypes and treatment history. We followed the standard quality control practices in scRNA-seq by performing quality control and subsequent filtering. For the purpose of this study, we selected 5/42 samples based on their higher median genes per cells. This was followed by inferring tumor cells using inferCNV **(Tickle et al, 2019)** and identification non-tumor cell types including T cells, B lymphocytes, myeloid cells, neutrophils, mast cells, and follicular dendritic cells and stromal cell types (fibroblasts and endothelial cells). Re-clustering of each main cluster of non-malignant cells, identification of cluster markers, manual annotation and cross-referencing with external signatures **(Azizi et al, 2018; Cahoy et al, 2008; Lein et al, 2007; Olah et al, 2020; Zilionis et al, 2019)** yielded cell type labels for subpopulations. One of the benefits of using KINOMO (or factorization approaches in general) in scRNA-seq is to highlight intra- and inter-tumoral heterogeneity. We start our discussion with the *intra-tumoral heterogeneity* analysis. Conventional NMF approaches share some coherent challenges with their signed dimensional reduction counterparts. For example, identifying the factor or factor which could ‘partition’ the scRNA-seq data in a manner which highlights the intra-tumor heterogeneity, remains a challenging problem. For this purpose, KINOMO uses a consensus approach using iterative factor updating with a stopping criterion. We use a consensus approach based on **Brunet et al, 2004, Hutchins et al, 2008**, and **Frigyesi et al, 2008** methods. We then iteratively update the factor or factor until a convergence is achieved (discussed in details in ***Materials and Methods***).

For the purpose of illustration, using the consensus approach, we select the ‘best’ factor and the ‘2^nd^ best’ factor which could partition the scRNA-seq data. Here, factor is based on the cophenetic correlation coefficient to measure the stability of the partitions. General notion says that as the number of factors increase, the sample is more heterogenous. Thus, intra-tumoral heterogeneity is proportional to the number of factors as obtained for each sample. The ‘best’ factor defines the lowest factor required to obtain stable partition, whereas the ‘2^nd^ best’ factor is considered if ‘best’ factor cannot be used to obtain stable partition. For instance, the best factor in GSM4453576_P1 is 3 and the second best is 5 **(Figure 2a)**. It is also important to note that each of the factors is defined by a metagene signature which is unique. This metagene signature defines the gene expression program of a specific factor in an individual sample. As discussed in the ***Materials and Methods*** section, we identify the top 30/50/100/200/300 metagenes per factor **(Figure 6)**. For the purpose of illustration **Figure 2b** provides basis heatmaps for top 30 metagenes (based on integrated factorization weight). As an example, the factors in GSM4453576 have signature enriched in epithelial-to-mesenchymal transition (EMT) pathway genes *IFITM3, TMSB4X, SAT1, VIM, LGALS1* (Rank_1); reactive oxygen species pathway genes *PERP, TXN, ATP5MC3, NIPSNAP2, AKR1C2* (Rank_2), and myogenesis pathway genes *DSC2, TMPRSS11E, CSTB, KRT6A, KLF5* (Rank_3), respectively **(Figure 2b)**.

**Figure 2.**
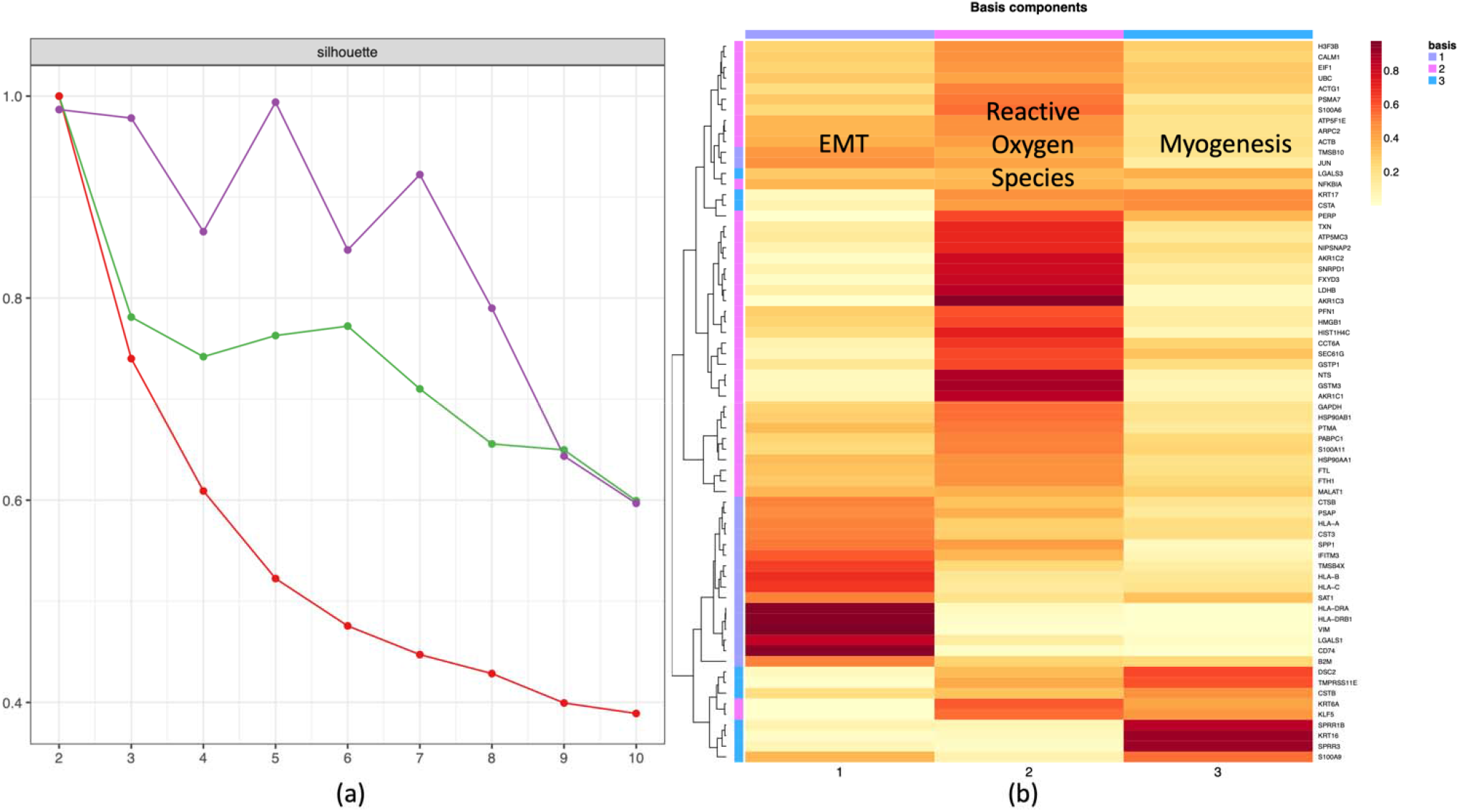
Intra-tumor heterogeneity in NSCLC (GSM4453576_P1), (a) Factor estimation identified ‘best’ factor = 3 and ‘2^nd^ best’ factor = 5. Here, the x-axis represents factor estimation (2–10) and the y-axis the silhouette score, (b) Basis heatmap using ‘best’ factor = 3 for top 30 metagenes reveals three distinct gene expression programs (EMT, reactive oxygen species and myogenesis). Here, the x-axis is the estimated basis components, y-axis is metagenes.

For the purpose of assessing the inter-tumor heterogeneity, we estimate the factor blocks (group of multiple factors) or ‘FBs’ by doing a simple co-correlation clustering using Spearman’s factor correlation for all factors across all samples **(Figure 3)** and perform Gene-set enrichment analysis (GSEA) (**Subramanian et al, 2005**) to estimate the corresponding pathways enriched for the factor blocks (FBs). Biologically, we are trying to identify 1) recurrently occurring modules across all samples, which signify how similar the gene expression programs across samples are, and 2) the level of inter-tumor heterogeneity by identifying unique FBs. For the five samples, we obtain 15 factors and based on the co-correlation analysis we identify three factor blocks, namely, FB1 (2 factors, GSM4453576_P1_R3_F1, GSM4453584_P9_R3_F2), FB2 (2 factors, GSM4453584_P9_R3_F1, GSM4453616_P41_R3_F2) and FB3 (8 factors, GSM4453576_P1_R3_F2, GSM4453578_P3_R3_F2, GSM4453578_P3_R3_F3, GSM4453592_P17_R3_F1, GSM4453592_P17_R3_F2, GSM4453592_P17_R3_F3, GSM4453616_P41_R3_F1, GSM4453616_P41_R3_F3). By performing a more granular analysis, we see that FB1 is enriched in gene signature for NF-kB, FB2 enriched in EMT, and FB3 enriched in cell cycle **(Figure 3)**. Interestingly, cell cycle is one of the biggest gene expression programs in cancer. Cell-cycle checkpoints are compromised in cancer cells to allow continuous cell division. This work is guiding and improving existing therapeutics and highlights opportunities to develop novel and combinatorial treatments in cancer treatments. These specifically include targeting replication stress tolerance mechanisms, the mitotic checkpoint, and proteins and processes involved in delaying or arresting cell-cycle progression **(Matthews et al, 2022)**. Interestingly, in recent years, inflammation has been established as a key inducer of EMT during the progression of cancer **(Mantovani et al, 2008)**. Modification of the TME during EMT occurs as a result of the activity of cytokines, such as IFN-γ, TGF-β and TNF-α which have been shown to induce EMT during cancer progression **(Ricciardi et al, 2015; Grivennikov et al, 2010)**. Likewise, spectrum of EMT states in NSCLC promises insights on cancer progression and drug resistance **(Karacosta et al, 2019)**. Likewise, NF-κB is a master regulator, not only of the physiologic and complicated process of lung morphogenesis but also of lung cancer pathogenesis and progression. It is known that NF-κB in NSCLC has a bidirectional contribution because, on one hand, it plays a crucial role in the immune response, whereas on the other hand, it promotes the inflammation that ignites the process of oncogenesis **(Aggarwal, 2004)**. Hence, activated NF-κB signaling may influence the progression of a lung tumor either positively or negatively **(DiDonato□et al, 2012)**. It is well documented that NF-κB is activated by a great variety of carcinogens, chemotherapeutics, cytokines, and radiation exposure **(Bharti□et al, 2012)**. In addition, it has been reported that NF-κB participates in and orchestrates many significant functions that tumors require, such as transformation, proliferation, infiltration, angiogenesis, and metastasis. Taken together, KINOMO is able to pick up essential gene expression programs cancer that could potentially lead to target-based precision therapies.

**Figure 3.**
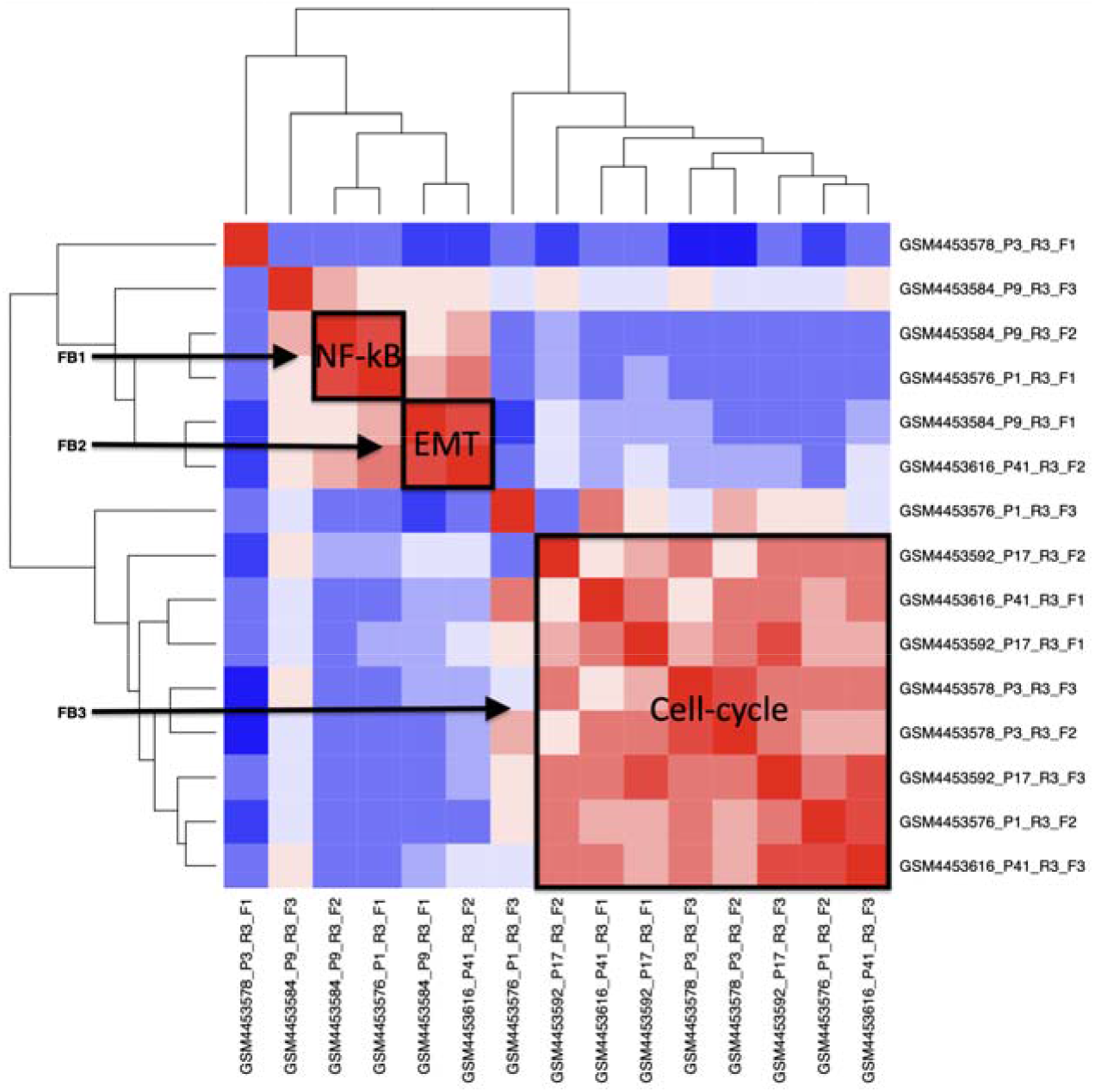
Factor block estimation for NSCLC data using co-correlation matrix of factors/factors across five samples. Three factor blocks can be assigned (FB1: enriched for NF-kB; FB2 enriched for EMT; and FB3 enriched for cell cycle).

## Discussion

In this manuscript we propose KINOMO (**K**ernel d**I**fferentiability correlation-based **NO**n-negative **M**atrix factorization algorithm using Kullback-Leibler divergence loss **O**ptimization), a factorization method that is robust to noise, explicit to error correction and also uses ‘prior’ biological knowledge for studying inter-tumoral heterogeneity. This prior knowledge in KINOMO is based on Gaussian process priors that are linked to the non-negative factors by a link function. More specifically, these Gaussian process priors are based on the multiple modalities of scRNA-seq data. This can also be combined with any existing NMF cost function that has a probabilistic interpretation, followed by using unconstrained optimization algorithm for computing the maximum a posteriori estimate.

Given an input data matrix, any existing NMF finds an approximation that is factorized into a product of lower-rank matrices, some of which are constrained to be non-negative. This is followed by optimizing the approximation error that is measured by a variety of divergences between the input and its approximation **(Kompass 2006; Dhillon et al, 2006; Cichocki et al, 2008)**, whereas the factorization can take a number of different forms **(Pascual-Montano et al, 2006; Ding et al, 2006; Ding et al, 2010)**. KINOMO uses a quadratic approximation approach to optimize the factorization error and uses a more generalized sequential quadratic approximation for KL divergence loss. This is achieved when *L* is a square loss and considering a sequential coordinate-wise descent (SCD) method proposed by **Franc et al, 2005** is used to solve a penalized non-negative least square for *H* while *W* is fixed. Another benefit of using SCD-based approach for KL divergence loss also deals with handling the time complexity, since it follows the Newton-Raphson like second order approach. Other NMF approaches like Scalar Block Coordinate Descent (sBCD) **(Li et al, 2012)** have used Bregman divergences to measure the quality of approximating a given matrix by the product of two lower matrices with non-negative indices. **Li et al, 2012** also define a new relationship connecting Bregman divergences (Frobenius norm) using Euclidean distance via Taylor series expansion. Further, a local updating rule is obtained by setting the gradient of the new objective function to zero with respect to each element of the two matrix factors. KINOMO uses a more generalized form of Bregman divergence, the Kullback-Leibler (KL) for approximation and *L*_*2,1*_ *norm loss* in place of Frobenius norm. *L*_*2,1*_ regularization optimizes the mean cost which is often used as a performance measurement. This is especially good since we want to keep the overall error small. The solution is more likely to be unique, wherein the non-sparseness of *L*_*2,1*_ improves KINOMO’s prediction performance. Finally, *L*_*2,1*_ is invariant under rotation, tends to shrink coefficients evenly and is useful since we have collinear and codependent features.

Most NMF methods are linear since each factorizing matrix appears only once in the approximation. However, such linearity assumption does not hold in case of extremely sparse data such as scRNA-seq, but rather the approximation is quadratic. Conventionally, quadratic approximations are more complex since these usually involve a higher-degree objective function with respect to the doubly occurring factorizing matrices **(Yang et al, 2012)**. KINOMO also uses quadratic approximation but is different from **Yang et al, 2012**, and uses a more generalized Lagrangian form of NMF, since conventional NMFs like **Yang et al, 2012** suffer from convergence problems. The novelty of KINOMO’s implementation of the quadratic approximation (which is convex in nature) is that it links it to the SCD approach, which is more similar to an Incremental Gradient Descent algorithm. The quadratic form of NMF as discussed by **Yang et al, 2012** considers using squared Euclidean distance-based divergences. KINOMO implements an efficient approximative second-order optimization based on multiplicative updates for faster convergence.

Random initialization is the benchmark used in the vast majority of NMF analysis. Probabilistic concepts are one of the useful approaches for NMF initialization. Moreover, random initialization is a common approach for NMF relying on a random selection of columns of the input matrix. These different random initializations can lead to different solution paths toward local minima, as finding the global minimum is challenging. This is somewhat counteracted by setting a random seed to guarantee reproducibility of a factorization model. However, in scRNA-seq, replicate models give similar errors of reconstruction, wherein many factors are robust, and some even share information among themselves. Compared to conventional NMF and other clustering methods, KINOMO has higher accuracy in predicting factors, thereby helps identifying hidden structures in scRNA-seq data across patients with varying genomic load. Finally, using conventional GSEA strategies, we observe that the estimated factor blocks are enriched in unique gene signatures and pathways. In conclusion, KINOMO provides an accurate approach to identifying recurrent features of cellular variability in single-cell RNA-seq data analyses of cancer tissues.

## Materials and Methods

### Methods used for benchmarking

To benchmark KINOME, we used hierarchical clustering **(Maimon et al, 2006)**, k-means **(Hartigan et al, 1979)**, Fuzzy C-Means **(Dunn 1973)**, SC3 **(Kiselev et al, 2017)**, TSCAN **(Zhicheng et al, 2016)**, SAFE **(Yuchen et al, 2019)**, FlowSOM **(Van Gassen et al, 2015)**, CIDR **(Lin et al, 2017)** and NMF **(Lee et al, 1999) (Table 2)**.

Hierarchical clustering is a general family of clustering algorithms for building nested clusters by merging or splitting them successively. This hierarchy of clusters is represented as a tree. The root of the tree is the unique cluster that gathers all the samples, the leaves being the clusters with only one sample. We use a division-based algorithm which initially starts with all observations in a single cluster, followed by dividing samples until each cluster only contain one observation and use Euclidean distance as the metric **(Maimon et al, 2006)**. The k-means algorithm clusters data by trying to separate samples in *n* groups of equal variances, minimizing a criterion known as the inertia or within-cluster sum-of-squares. k-means requires the number of clusters to be specified and scales well to large number of samples. It divides a set of *N* samples *X* into *K* disjoint clusters *C*, each described by the mean *µ*_*j*_(centroid) of the samples in the cluster. It aims to choose centroids that minimize the inertia, or within-cluster sum-of-squares criterion: 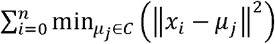 **(Hartigan et al, 1979)**.

Fuzzy C-Means is a soft-clustering approach, where each data point is assigned a likelihood or probability score to belong to that cluster **(Dunn 1973)**. It has four major steps, namely, i) fixing number of clusters, *c*; select a fuzziness parameter, *m* (generally 1.25 < m < 2); initialize partition matrix, 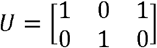; ii) calculate centroid, 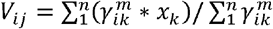; iii) update partition matrix, 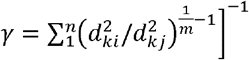 and iv) repeat steps i-iii until convergence. SC3 computes a consensus matrix using the cluster-based similarity partitioning algorithm (CSPA) **(Strehl et al, 2003)**. For each individual clustering result a binary similarity matrix is constructed from the corresponding cell labels: if two cells belong to the same cluster, their similarity is 1, otherwise 0. This is followed by calculating a consensus matrix by averaging all similarity matrices of individual clustering. The resulting consensus matrix is clustered using hierarchical clustering with complete agglomeration and the clusters are inferred at the *k* level of hierarchy, where *k* is defined by a user **(Kiselev et al, 2017)**.

TSCAN (Tools for Single Cell Analysis) is a tool for differential analysis of scRNA-seq data, which uses a cluster-based minimum spanning tree (MST) method for the pseudo-temporal ordering of cells. If we consider a representative sample of *N* cells drawn from a heterogeneous cell population and suppose that the transcriptome *Y*_*i*_ of each cell *i* _∈_ *{1, 2, …, N}* has been profiled using scRNA-seq, we assume that, *Y*_*i*_ is a *G*-dimensional vector consisting of gene expression measurements for *G* genes. Further, assuming that *Y*_*i*_ is appropriately transformed and normalized across cells, the single-cell ordering problem or pseudo-time reconstruction, is to place cells in an order based on the gradual transition of *Y*_*i*_. TSCAN orders cells in three steps, namely, i) cells with similar gene expression profiles are grouped into clusters, ii) a minimum spanning tree (MST) is constructed to connect all cluster centers, and iii) cells are projected to the tree backbone to determine their pseudo-time and order **(Zhicheng et al, 2016)**. Single-cell Aggregated (From Ensemble) (SAFE) clustering leverages hypergraph partitioning methods to ensemble results from multiple individual clustering methods. It embeds four clustering methods: SC3, Seurat, t-SNE + k-means and CIDR. SAFE-clustering takes an expression matrix as input, followed by converting the fragments/reads per kilobase per million mapped reads (FPKM/RPKM) data are converted into transcripts per million (TPM) and UMI counts into counts per million mapped reads (CPM). This is followed by conventional downstream analysis **(Yuchen et al, 2019)**.

FlowSOM is a powerful clustering algorithm that builds self-organizing maps to provide an overview of marker expression on all cells and reveal cell subsets that could be overlooked with manual gating. It has been shown to produce results rapidly and automatically groups cell clusters into higher order meta-clusters **(Van Gassen et al, 2015)**. Clustering through Imputation and Dimensionality Reduction (CIDR) algorithm has five steps: (i) Identification of dropout candidates, (ii) estimation of the relationship between dropout rate and gene expression levels, (iii) calculation of dissimilarity between the imputed gene expression profiles for every pair of single cells, (iv) PCoA using the CIDR dissimilarity matrix, and (v) clustering using the first few principal coordinates **(Lin et al, 2017)**.

Finally, we also established whether components inferred by simple matrix factorizations would align with GEPs in scRNA-seq data. For each of the 5 samples, we generated 10 replicates, each at three different ‘signal to noise’ ratios, in order to determine how matrix factorization accuracy varies with noise level. We ran each method 100 times and assigned the components in each run to their most correlated ground-truth program. To evaluate the clustering performance of the different algorithm, we also use adjusted Rand index (ARI) which ranges from −1 to +1 **(Rand 1971)**.

## Author Contributions

S.T. and B.I. conceived the study. B.I. provided overall supervision. Y.W. and J.B. contributed to the analysis. R.R. and E.A. contributed to developing the model. S.T., E.A. and B.I. wrote the manuscript.

## Acknowledgements

is supported by National Institute of Health (NIH) National Cancer Institute (NCI) grants K08CA222663, R37CA258829, R21CA263381 (with E.A.), U54CA225088 and U54AG076040, an American Cancer Society Research Scholar Grant, a Burroughs Wellcome Fund Career Award for Medical Scientists, a Velocity Fellows Award, the Louis V. Gerstner, Jr. Scholars Program, a V Foundation Scholar Award, a Columbia University RISE award (with E.A.), and the Tara Miller Young Investigator Award by the Melanoma Research Alliance.

